# PCR procedures to amplify GC-rich DNA sequences of *Mycobacterium bovis*

**DOI:** 10.1101/2020.02.18.953695

**Authors:** Nadia Assal, Min Lin

**Affiliations:** Ottawa Laboratory Fallowfield, Canadian Food Inspection Agency, Ottawa, Ontario, Canada; Department of Biochemistry, Microbiology and Immunology, University of Ottawa, Ottawa, Ontario, Canada

## Abstract

Amplification of high GC content genes by PCR is a major challenge during the creation of recombinant GC-rich DNA constructs. This may be due to the difficulty in DNA denaturation or the possibility of forming secondary structures from DNA templates. Tools have been described to address the technical problems associated with the amplification of shorter sequences (<1000 bp). However, obstacles of synthesizing larger-sized GC-rich sequences by PCR continue to exist. This study aims to investigate the amplification of long and high GC content genes by PCR from the *Mycobacterium bovis*, a genome with GC content >60%, in comparison to amplifying a gene from the *Listeria monocytogenes* genome, a genome with a 37.8% GC content. Three PCR protocols were designed and experimented at various conditions with two *M. bovis* genes, *Mb0129*, a large gene of 1794 bp with 77.5% GC content, *mpb83*, a smaller gene of 663 bp in length with moderate GC content of 63%, together with *LMHCC_RS00060*, a large *L.monocytogenes* gene of 1617 bp with a lower GC content of 41.53%. The result demonstrated the superiority of the 2-step PCR protocol over other protocols in PCR amplification of *Mb0129* when specific high fidelity DNA polymerases were used in the presence of an enhancer. The study highlighted the importance of manipulating the cycling conditions to perform the annealing and extension steps at higher temperatures for a successful PCR amplification of a large GC-rich DNA template. The PCR protocols developed in this study can be valuable tools for the amplification of long GC-rich DNA sequences for various downstream applications.

## Introduction

The development of the polymerase chain reaction (PCR) amplification of DNA has been considered a revolution in molecular biology since its discovery [1]. It is an indispensable tool in both medical and biological research fields. Applications of PCR include the diagnosis of infectious diseases, molecular genetic analysis, and the creation of recombinant DNA constructs for protein expression purposes [2,3]. Despite being a straightforward technique, amplification of target DNA sequences of high GC content (>60%) can be an obstacle to a successful PCR. GC-rich sequences are present in important regulatory elements of human DNA including promoters and enhancers [4], in addition to some bacterial genomes characterized by high GC content such as Mycobacteria and Pseudomonas [5,6]. Cloning genes of high GC content by PCR using long primers can pose an extra challenge to the amplification process because of the high melting temperature (*Tm*), added to the possibility of forming secondary structures such as hair pin, self and cross dimer formation [7].

Numerous studies aiming to overcome problems associated with PCR amplification of high GC sequences are documented in the literature. Special consideration is given to the primers, enhancer solutions and cyclic conditions. It is recommended to keep the length of the primers between 15 to 30 nucleotide residues (bases) and to have the *Tm* between 52-58 °C. Moreover, di-nucleotide repeats (e.g., GCGCGCGCGC) should be avoided to prevent hair pin structure formation [7]. Designing such primers can be achieved using a commonly used program, Primer3 (http://primer3.ut.ee/). The addition of enhancers can modify the melting characters of the double-stranded DNA helices allowing easy separation of the strands. The most commonly described in the literature are betaine, dimethyl sulfoxide (DMSO) and formamide. Betaine is an amino acid analog that decreases the energy required for DNA strands denaturation. DMSO interferes with the hydrogen bond formation preventing inter- and intrastrand reannealing. However, DMSO has the drawback of reducing the activity of Taq polymerase making it is important to consider maintaining a balance between template accessibility and the polymerase activity [8–11]. Formamide increase PCR specificity when working with GC-rich targets [12]. Modification of cyclic conditions is also considered; this includes optimizing the annealing temperature, applying Hot start PCR, and using nested PCR and touchdown PCR [13]. Touchdown PCR (TD-PCR) uses an annealing temperature higher than the expected annealing temperature. This higher temperature is decreased 1-2°C every cycle usually for ten cycles until the desired annealing temperature is reached. TD-PCR makes benefit from the exponential nature of PCR reactions giving an advantage of 2-fold per cycle for the production of the desired product [13,14]. The use of TD-PCR has been described to overcome multiple issues including difficulties in the amplification of high GC content targets [15].

It is noted that even with adopting the previously mentioned approaches, amplification challenges still exist especially when using GC-rich templates, lengthy primers and aiming to amplify longer products. Moreover, most of the published literature focuses on the amplification of genes less than 1 kilobase and optimization of the conditions relating to a single PCR product [5,8,9,16,17], however, they do not address working with multiple lengthy GC-rich targets (>1kb). In the context of another project aiming for amplification and cloning of more than 100 GC-rich ORFs from the *Mycobacterium bovis* genome, a standard procedure that enables amplification of multiple targets simultaneously was required without the need to optimize individual conditions for each target. This provoked us to design this study that tested and compared the PCR amplification of one “difficult” target as a model, *Mb0129* from *M. bovis*, a large-sized gene 1794 bp and a GC content 77.5%, and two relatively “easier” targets, *mpb83 (Mb2898)* from *M. bovis*, a 663 bp gene and 63% GC content, and *LMHCC_RS00060* from *Listeria monocytogenes*, a 1617 bp gene and 40% GC. Three PCR protocols were designed and experimented with each of the three genes using five different polymerase enzymes in the presence or absence of enhancers.

## Materials and Methods

### Genomic DNA

*M. bovis* genomic DNA was extracted using QIAGEN DNeasy blood and tissue kit according to the manufacturer’s instructions for Gram-positive bacteria. *L. monocytogenes* HCC23 serotype 4a DNA was extracted using DNAzol. Briefly, *L. monocytogenes* cells were cultured in BHI broth, the cell pellet was resuspended in TE buffer digested by lysozyme, followed by the addition of DNAzol and ethanol. The reaction was centrifuged at 13000 rpm for 1 min, the supernatant was discarded and the DNA pellet was dissolved in NaOH. Five ng/μl of genomic DNA was used in each reaction.

### Primers

For the purpose of *in vivo* cloning these genes into a single vector (pIVEX2.3d) (Biotechrabbit, Germany), primers were designed to consist of a tail of 26 bp in forward primer (FP) and 25 bp in reverse primer (RP) homologous to the ends of the cut vector in addition to 20 bp or more specific to the target genes, Table 1. Sequences of homology allowed homologous recombination between the vector and the target sequences in *E.coli* [18].

**Table 1.**
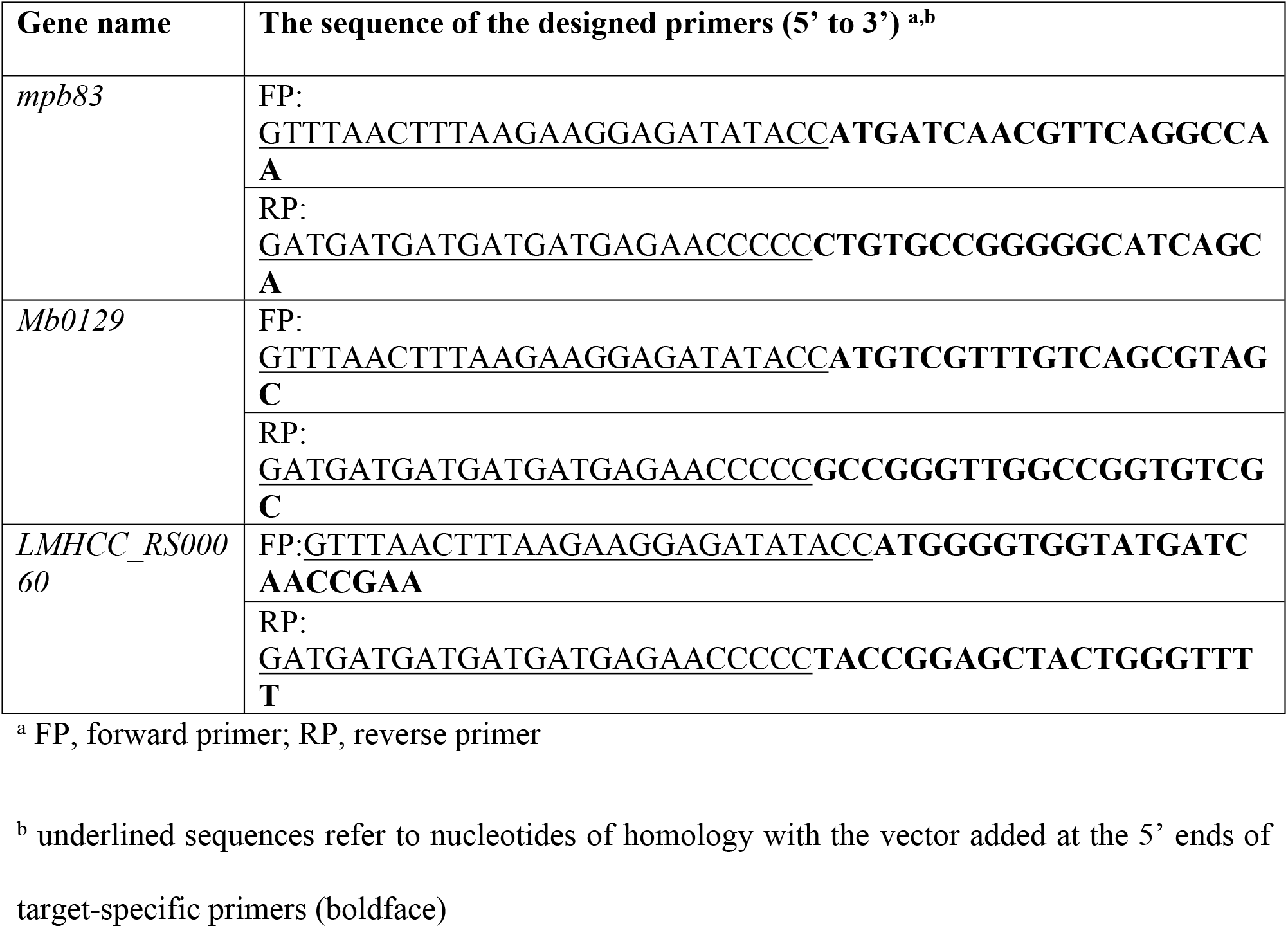
The designed primers for the reactions.

### Amplification enzymes

Amplification of the genes was done using high fidelity polymerase enzymes including Phusion® High-Fidelity DNA Polymerase (New England Biolabs, NEB Cat# M0530), Q5® High-Fidelity DNA Polymerases (NEB, Cat# M0491), Platinum™ SuperFi™ DNA Polymerase (Invitrogen, **Cat**# 12351010), PrimeSTAR GXL DNA Polymerase (Takara Bio, Cat# R050Q) and Taq polymerase (home-made).

### Master mixes

Enhancers specified in the polymerase kit were used if supplied, otherwise 2-5% DMSO (Sigma, Cat# D9170) were used instead. Master mixes were prepared following the manufacturer’s instruction as two preparations for each candidate; one with an enhancer and one without an enhancer. Those with the enhancer had either the maximum kit-supplied enhancer (10 μl per 50 μl reaction) or 5% DMSO (0.7M) for both *mbp83* and *Mb0129*. The *LMHCC_RS00060* reactions with the enhancer contained either 50% of kit-supplied enhancer or 2.5% DMSO (0.35M). The primers were added at 0.5 μM final concentration, and the dNTPs were at 0.2mM final concentration. The genomic DNA was used at 5ng/reaction. All experiments were done in a 50 μl final reaction volume.

### Annealing temperature calculation

Annealing temperatures for the 3-step PCR reactions were used as specified by the manufacturer recommendations, 60°C for PrimeSTAR GXL DNA Polymerase, or using the *T*m calculator on the manufacturers’ websites (http://tmcalculator.neb.com/#!/main) for Phusion and Q5 polymerases and (www.thermofisher.com/tmcalculator) for Platinum™ SuperFi polymerase.

### Thermal cycle conditions

Three protocols were used to assess the productivity of each enzyme on its own. Amplification using 3-step (3St), 2-step (2St) and a 3Step with touchdown (TD) protocols were attempted following the manufacturer’s recommendations for the different enzymes. The protocols for the 3St and 2St are represented in Table 2A, and the TD protocol is represented in Table 2B.

**Table 2A.** 3St and 2St protocol for the different enzymes used.

| Protocol |  | 3St |  |  |  |  |  | 2St |  |  |  |  |  |
| --- | --- | --- | --- | --- | --- | --- | --- | --- | --- | --- | --- | --- | --- |
| Enzyme |  | Phusion, Q5, SuperFi<sup>b</sup> |  | GXL<sup>b</sup> |  | Taq |  | Phusion, Q5, superFi |  | GXL |  | Taq |  |
| Step |  | Temp<sup>a</sup> | Time | Temp<sup>a</sup> | Time | Temp | Time | Temp<sup>a</sup> | Time | Temp<sup>a</sup> | Time | Temp<sup>a</sup> | Time |
| Initial denaturation |  | 98 | 30s | - | - | 95 | 2m30s | 98 | 30s | - | - | 95 | 2m30s |
| 35 cycles | denature | 98 | 10s | 98 | 15s | 95 | 20s | 98 | 10s | 98 | 15s | 95 | 20s |
|  | anneal | 63 | 20s | 60 | 15s | 60 | 20s | - | - | - | - | - | - |
|  | extend | 72 | 1m:30s | 68 | 2m | 72 | 2m30s | 72 | 1m:30s | 68 | 2m | 72 | 2m30s |
| Final extension |  | 72 | 5m | - | - | 72 | 5m | 72 | 5m | - | - | 72 | 5m |
| Hold |  | 4 | ∞ | 4 | ∞ | 4 | ∞ | 4 | ∞ | 4 | ∞ | 4 | ∞ |

<sup>a</sup>Temp refers to temperatures in °C
<sup>b</sup>“SuperFi” stands for Platinum™ SuperFi™ DNA Polymerase and “GXL” for PrimeSTAR GXL
DNA Polymerase

**Table 2B.** TD protocol for the different enzymes used.

| Step \ Enzyme |  | Phusion, Q5, SuperFi<sup>b</sup> |  | GXL<sup>b</sup> |  | Taq |  |
| --- | --- | --- | --- | --- | --- | --- | --- |
|  |  | Temp<sup>a</sup> | Time | Temp<sup>a</sup> | Time | Temp<sup>a</sup> | Time |
| Initial denaturation |  | 98 | 30s | - | - | 95 | 2m30s |
| 10 cycles | denature | 98 | 10s | 98 | 15s | 95 | 20s |
|  | anneal<sup>c</sup> | 72 (-1.6/ cycle) | 20s | 72 (-1.2/ cycle) | 15s | 72 (-1.6/ cycle) | 20s |
|  | extend | 72 | 1m30s | 68 | 2m | 72 | 2m30s |
| 30 cycles | denature | 98 | 10s | 98 | 15s | 95 | 20s |
|  | anneal | 55 | 20s | 60 | 15s | 55 | 20s |
|  | extend | 72 | 1m30s | 68 | 2m | 72 | 2m30s |
| Final extension |  | 72 | 5m | - | - | 72 | 5m |
| Hold |  | 4 | ∞ | 4 | ∞ | 4 | ∞ |

<sup>a,b</sup>as described in Table 2A
° For the TD PCR the annealing temperature decreases by 1.2 or 1.6 °C per cycle for the first ten cycles until it reaches the desired annealing temperature.

## Results and Discussion

Our group has recently experienced difficulties in our attempts to amplify various ORF targets in the *M. bovis* genome by PCR for the purpose of cloning. Most of the ORFs targets are more than 1 kb in size and have a GC content of more than 65% which presumably hindered PCR amplification of the *M. bovis* ORF targets with DNA polymerases under a conventional PCR cycling. Unsuccessful attempts were faced with Taq polymerase, One*Taq* DNA Polymerase (NEB, M0480S), Platinum™ Pfx DNA Polymerase (Invitrogen, 11708039), Expand Long Template PCR System (an enzyme mix that contains thermostable Taq DNA Polymerase and a thermostable DNA polymerase with proofreading activity, Roche, 11681834001). This observation prompted us to establish a reliable PCR procedure that not only can be used for cloning the *M. bovis* sequences but also can be applied to the amplification other targets of high GC content. Here we evaluated and compared several PCR protocols for the amplification of three selected ORFs of high, moderate and low GC content. These ORFs are *Mb0129* (a gene from *M. bovis* composed of 1794 bp with a GC content of 77.5%), *mpb83* (a smaller gene of 663 bp with a GC content of 63.0% from *M. bovis*), and *LMHCC_RS00060* (a large gene of 1617 bp with a GC content 41.5% from *L. monocytogenes*). Four high fidelity enzymes were chosen for use in the PCR protocols, based on their ability to amplify high GC-content genes as indicated by the suppliers, in addition to the widely used Taq polymerase. PCR amplification of the three selected targets was attempted with each of the 5 enzymes described in the Materials and Methods. Three PCR protocols, 3St, 2St, and TD protocols, were designed as per the instructions of the enzyme manufacturers or as described by Don et al. [15] and tested with each of the 5 selected DNA polymerases. Given that enhancers such as DMSO have been recognized for having the ability to improve the PCR performance [8,9], we also investigated the effect of the enhancers on PCR amplification of the target genes containing a high GC-content. A summary of all results is illustrated in Table 3.

**Table 3.** Summary of the PCR results from the amplification of three target genes by using 5 different DNA polymerases in the three protocols.

| Enzyme | Protocol | *mpb83* |  | *Mb0129* |  | *LMHCC_RS00060* |  |
| --- | --- | --- | --- | --- | --- | --- | --- |
|  |  | Enhancer added <sup>a</sup> |  | Enhancer added <sup>a</sup> |  | Enhancer added <sup>a</sup> |  |
|  |  | N | Y | N | Y | N | Y |
| **Taq** | **3 St** | - | + | - | - | - | + |
|  | **2 St** | - | + | - | - | - | + |
|  | **TD** | - | + | - | - | - | + |
| **Q5** | **3 St** | + | + | - | + | + | - |
|  | **2 St** | + | + | - | + | + | - |
|  | **TD** | + | + | - | - | + | + |
| **Platinum SuperFi** | **3 St** | + | + | - | + | + | + |
|  | **2 St** | + | + | - | + | + | - |
|  | **TD** | + | + | - | - | + | + |
| **Prime STAR GXL** | **3 St** | + | + | - | + | + | + |
|  | **2 St** | + | + | - | + | + | + |
|  | **TD** | + | + | - | - | + | + |
| **Phusion** | **3 St** | + | + | - | - | + | weak |
|  | **2 St** | + | + | - | weak | + | - |
|  | **TD** | + | + | - | - | + | weak |

<sup>a</sup> N refers to reactions done without the added enhancer, Y refers to those with the enhancer added
+ refers to a successful amplicon production, - refers to no amplicon production

### PCR amplification using Taq DNA polymerase

Taq is the most commonly used DNA polymerase in PCR for amplification of a target DNA in laboratories. With Taq DNA polymerase, three PCR protocols (Materials and methods) were designed and evaluated for amplifying selected ORFs of low, moderate and high GC contents. Both *mpb83* (63% GC) and *LMHCC_RS00060* (41.5% GC) were successfully generated by employing the three protocols in the presence of DMSO (an enhancer) at 5% and 2%, respectively. For *Mb0129* (77.5% GC), all of the three protocols failed to produce the amplicons in the presence of 5% DMSO, Fig 1A. Without DMSO, Taq failed to synthesize the PCR products for all three gene targets *mpb83* and *LMHCC_RS00060* and *Mb0129*, Fig 1B, suggesting that an enhancer is required for the PCR reactions. One benefit of using an enhancer in PCR is believed to prevent the formation of secondary structures in long primers. These results revealed the difficulty of amplifying a larger gene, especially a target of more than 1 kb in size and higher GC content (> 65%) by Taq. The touchdown protocol drastically improved the yield of both *mpb83* and *LMHCC_RS00060* amplification; however, it still failed to produce the *Mb0129* amplicon, Fig 1A.

**Fig 1.**
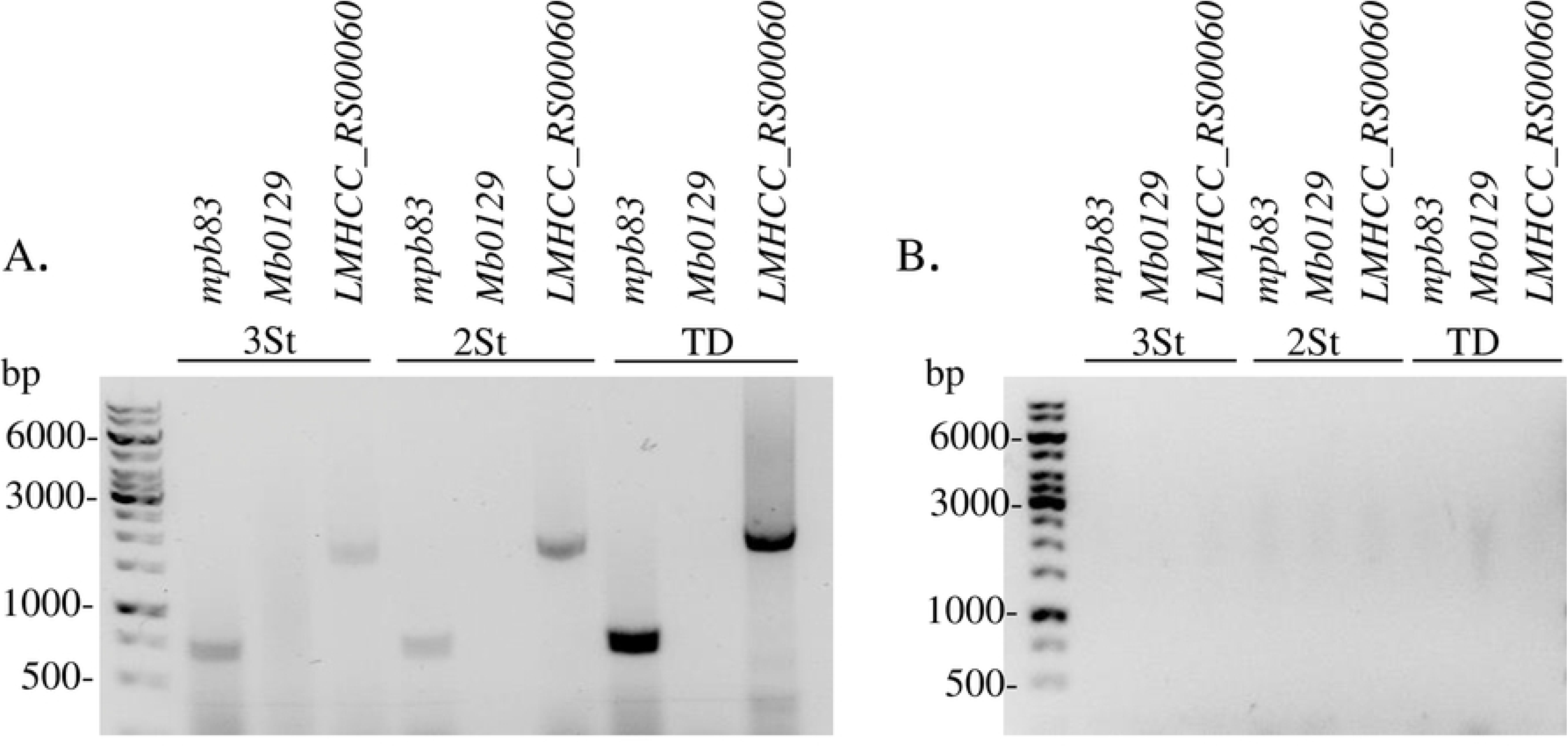
Amplification results of three target genes using Taq polymerase. (A) Attempted amplification of the three DNA targets (*mpb83*, *Mb012,* and *LMHCC_RS00060*) sequences by Taq in the presence of DMSO using 3St, 2St and TD protocols; *mpb83* and *LMHCC_RS00060* successfully showed amplicons using the three protocols. *Mb012* failed to show an amplicon. (B) Attempts of amplification of the three DNA target sequences in the absence of DMSO using 3St, 2St and TD protocols showing no amplicon. *mpb83* with added enhancer in 3St PCR is used as a positive control. *Mb0129* with no enhancer in 3St PCR is used as a negative control.

### The superiority of Q5 DNA polymerase over Phusion in the synthesis of *Mb0129*

Phusion and Q5 DNA polymerases were selected for use in the PCR protocols based on their high fidelity and suitability for amplification of high GC-content DNA templates and were used in the three protocols. Both enzymes were able to generate the *mpb83* amplicon in all the reactions of three PCR protocols with and without the presence of DMSO, Fig 2A. Despite the anticipated capacity of Phusion to amplify higher GC-content DNA sequences, two of the three protocols (3St and TD) failed to generate a detectable PCR product from the *Mb0129* template with this enzyme, while a 2St PCR reaction only weakly generated the *Mb0129* amplicon in the presence of DMSO. In contrast to Phusion, Q5 allowed the synthesis of *Mb0129* in good yield from the 2St reaction with the addition of the enhancer and produced a much lower yield of the amplicon in the 3St PCR reaction, Fig 2B. The low GC-content gene, *LMHCC_RS00060*, was successfully amplified by using the three protocols with either Phusion or Q5. PCR amplification of this gene appeared to be better without adding the enhancer DMSO in the reaction, Fig 2C. This may be attributed to the relatively low GC content of *LMHCC_RS00060*. These results demonstrated that Phusion worked well for the amplification of a shorter target sequence of moderate GC content in the presence of DMSO but not for a longer GC-rich sequence. Q5 DNA polymerase, when used in the 2St PCR protocol, was superior over Phusion in amplifying a longer DNA sequence of high GC-content.

**Fig 2.**
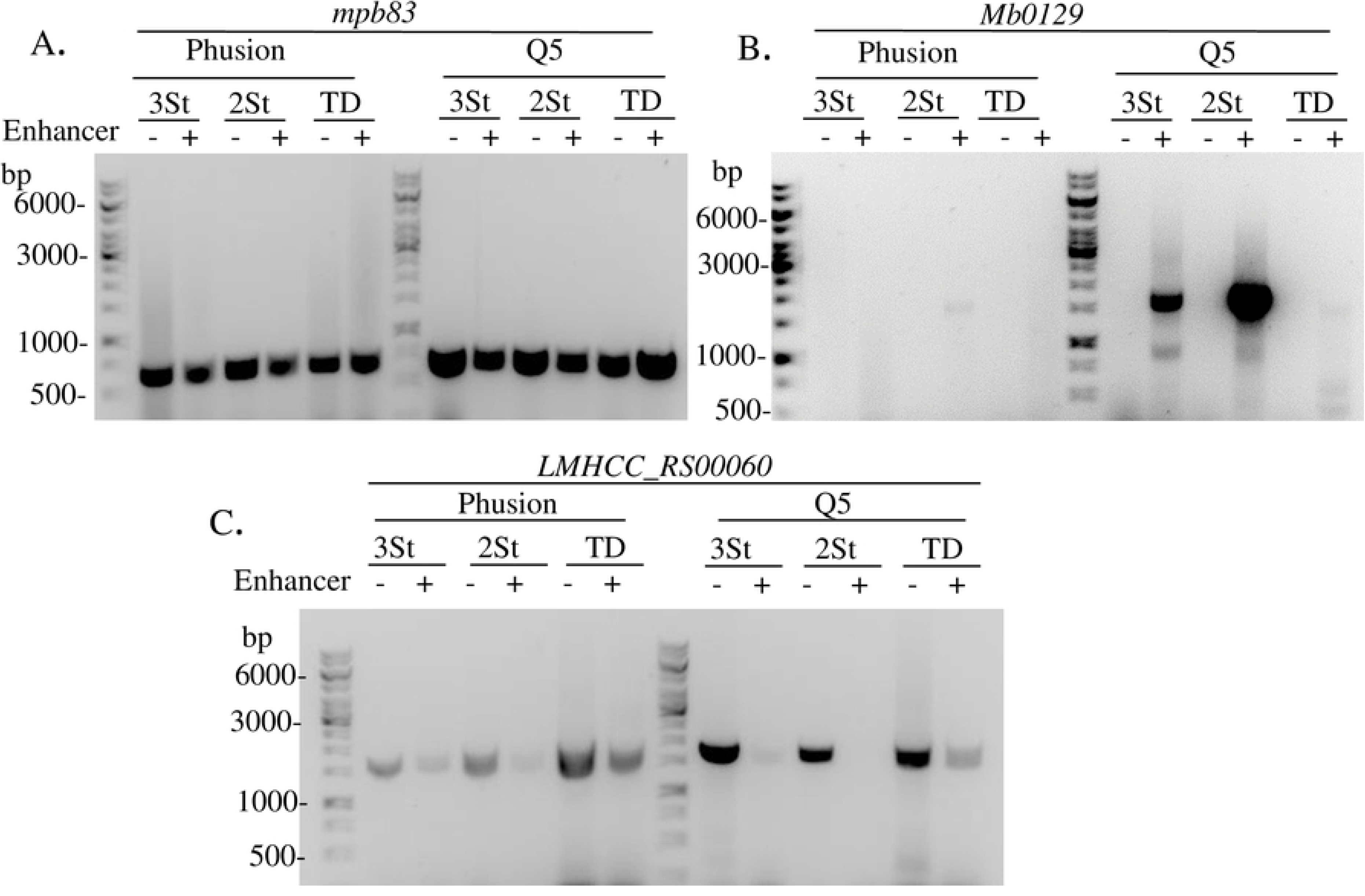
Results for attempted amplification of the three target genes using Phusion and Q5 polymerase. (A) *mpb83* is readily amplified using both Phusion and Q5 polymerase using the three different protocols. (B) *Mb0129* is only amplified using the Q5 enzyme using the 2St and 3St protocols in the presence of the enhancer. (C) *LMHCC_RS00060* shows amplicons with both enzymes in the three protocols with no need for the enhancer. Products are shown in the presence (+) and absence (−) of the enhancer. *mpb83* with added enhancer in 3St PCR is used as a positive control. *Mb0129* with no enhancer in 3St PCR is used as a negative control.

### The successful synthesis of *Mb0129* by using PrimeSTAR GXL and Platinum SuperFi polymerases

All the three PCR protocols with PrimeSTAR GXL DNA polymerase were able to amplify *mpb83* in all attempts. With this enzyme, both the 3St reaction and the 2St reaction produced a good yield of the *Mb0129* amplicon in the presence of 5% DMSO, Fig 3A. Similarly, Platinum SuperFi polymerase was capable of amplifying *mpb83* by using all three PCR protocols and behaved similarly in amplifying *Mb0129* via the 3St and 2St reactions in the presence of the enhancer, Fig 3B. *LMHCC_RS00060* was readily amplified by both enzymes with no requirement for the enhancer, Fig 3C, and 3D.

**Fig 3.**
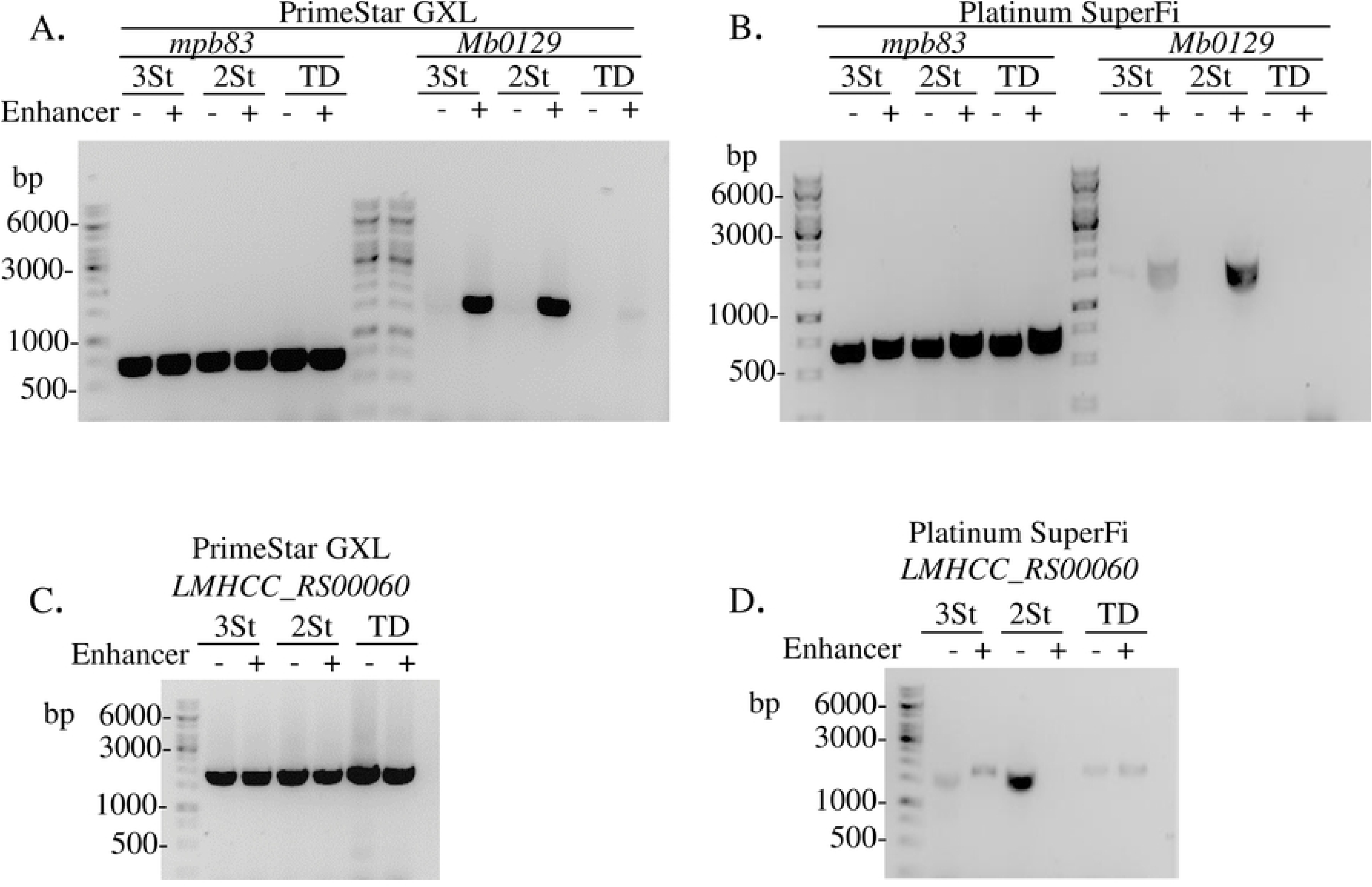
Results for attempted amplification of the three target genes using PrimeSTAR GXL and Platinum SuperFi polymerase in the three protocols. (A) With PrimeSTAR GXL polymerase, *mpb83* is readily amplified by using the three different protocols whereas *Mb0129* is only amplified with the 2St and 3St protocols in the presence of the enhancer. (B) Amplification of *mpb83* and *Mb0129* with Platinum SuperFi polymerase show similar results as in panel A. *LMHCC_RS00060 are successfully amplified using* PrimeSTAR GXL polymerase (C) and Platinum SuperFi polymerase (D) without the need for the enhancer. The presence (+) and absence (−) of the enhancer in PCR reactions are indicated. *mpb83* with added enhancer in 3St PCR is used as a positive control. *Mb0129* with no enhancer in 3St PCR is used as a negative control.

### Comparison of the three PCR protocols

Examination of the results obtained with the 3St, 2St, and TD PCR protocols revealed that *mpb83*, the short gene (<1kb) with 63% GC content, and *LMHCC_RS00060*, the long gene (>1.5 kb) with 40% GC were easily amplified in most of the reactions and protocols, Table 3. On the other hand, *Mb0129,* the long DNA target (> 1.5 kb) with a high GC-content (77.5%) behaved differently. *Mb0129* amplification was achieved by both the 2St and 3St protocols, but not the TD protocol. The success of implementing the 2St or 3St PCR protocols is dependent on the use of a selected high fidelity DNA polymerase such as Q5, Platinum SuperFi, and PrimeSTAR GXL and of the enhancer. Apparently, the PCR protocol involved in the 2St reaction performed better than other protocols. The 3St PCR protocol performed similarly compared to the 2St protocol in some cases but generated a low yield for the amplification of *Mb0129* using Q5 and Platinum SuperFi polymerases.

Touchdown PCR has been widely recommended for the amplification of high GC-content genes. This method was also suggested for use in cases where the determination of primers’ melting temperatures is difficult [13,15]. The advantage of the TD PCR procedure is in exploring the exponential nature of PCR. Given that primer annealing and extension are very critical steps for PCR, starting by annealing at higher temperatures then decreasing the annealing temperature gradually gives a two-fold advantage per cycle towards the correct product synthesis [14]. As opposed to the expectation, the TD protocol failed to synthesize the larger GC-rich gene *Mb0129* while using several selected enzymes, although it was able to amplify the small gene *mpb83* of moderate GC-content and the large gene *LMHCC_RS00060* of low GC content.

The success in amplifying the *Mb0129* gene by the 2St PCR reaction but not the TD protocol indicated that the PCR reaction, including specific primer annealing and extension, works better at higher temperatures for a long GC-rich DNA template. In the TD protocol, the final annealing temperature at 55°C, lower than that used in the 2St or 3St protocol, may be a factor responsible for the observed difference in performance between the protocols. It should not be surprising that the temperature played a critical role in the PCR protocols for the amplification of the *M. bovis* ORFs especially with the use of lengthy primers. Primers were designed to clone the target ORFs into a plasmid vector, in which a stretch of nucleotides derived from the vector sequence was added at the 5’ end of gene-specific primers, Table 1. These long primers, with extra non-gene specific sequences, exhibit high melting temperatures (≥72°C) thus giving an advantage to the 2St PCR. In addition, it is highly possible that the high GC content of the *M. bovis* genome forms a hairpin or other secondary structures at the PCR annealing stage and thus interferes with the function of the polymerase by hindering product synthesis; another factor that favours a higher temperature in the reaction.

The results obtained with the combination of three factors showed a better performance with the 2St PCR. These factors are the GC-rich DNA template, the long DNA target (>1kb) and the use of long primers. We aimed to investigate the role of the length of the primers in these reactions. Takara PrimeSTAR GXL enzyme, was utilized to amplify a 1737 bp (fragment of *Mb0287c*), of 78% GC content from the genome of *M. bovis*. Primers were designed using the “Primer 3” software to have a short length of 20 nucleotides. The primers had a calculated melting temperature of 63°C. Results show the successful amplification of the target using 2St PCR with annealing and extension at 68 °C. Conversely, the 3St PCR with annealing at 63 °C did not produce an amplicon, S1Fig. It is clear that the primers with shorter lengths did not result in the success of the 3St PCR to amplify the long (1.7kb), GC-rich gene. This indicates that the two factors, a long target and GC-rich template, may be the key players that require higher temperature and thus favour the 2St PCR for a successful amplification.

The ramp speed of the thermal cycler may play a role in amplifying difficult targets by decreasing the ramp speed. Attempts were made to amplify a total of 7 genes (having lengths between 1617-2166 bp and GC content between 60-77.6%, S1 Table, by using PrimeSTAR GXL enzyme in a 2St PCR. The thermal cycle at a full ramp speed (6 degrees/s) and a lower speed (2 degrees/s) were explored. Results show the successful amplification of the 7 targets with the lower ramp speed, in contrast to only a single target amplified using a full ramp speed, S2 Fig. These results agree with the findings of Frey at al. [17] using a slow ramp speed for amplification of GC-rich sequences. Frey et al. [17], however, used an additional step in which they added 7’-deaza-2’-deoxyguanosine, a dGTP analog which disrupts hydrogen bonds and decrease the annealing temperatures. This step would disrupt the gene structure for the downstream cloning application and thus was not implemented in our experiments.A final version of a ready to work protocol is found on dx.doi.org/10.17504/protocols.io.bbxyippw.

Amplifying GC-rich templates and long targets can be a real obstacle in PCR and may need a lengthy time for optimization. In this study, we found that selected polymerase enzymes tailored for this purpose like PrimeSTAR GXL enzyme, Platinum SuperFi and Q5 with the use of the enhancers are indispensable for amplifying a GC-rich gene (>60%) of more than 1kb in length. Adjusting the cyclic conditions to allow annealing and extension at high temperatures is offered by the 2St PCR and creates a favourable environment to amplify these targets. Slowing down the ramp speed of the thermal cycler is an additional solution to amplify difficult targets. Implementation of these recommendations can prevent the secondary structure formation, leading to a successful PCR amplification.

## Acknowledgment

We thank Dr. Dele Ogunremi, Dr. Mingsong Kang and Dr, Ray Theoret at the Canadian Food Inspection Agency for their insightful comments. We also thank Dr. Olga Andrievskaia for supplying us with *M. bovis* DNA and for her great suggestions for our experiments.

## Supporting information

**S1 Fig. Primer design and amplification results for *Mb0287c* amplification**.

A. The sequence of the primers (5’ to 3’) designed for the amplification of a 1737 bp sequence (fragment of *Mb0287c*). B. An agarose gel electrophoresis picture of the amplification results showing successful amplification of *Mb0287c* with the 2St PCR, lane1, whereas no product of *Mb0287c* with 3St PCR, lane 2. Lanes 3 and 4 show the positive control (*mpb83*) amplified while using an enhancer in 2St PCR and 3St PCR respectively. Lanes 5 and 6 show the negative control (*Mb0129*), with no enhancer added, and amplified using 2St PCR and 3St PCR respectively.

**S2 Fig. Comparison of the effects of modification of the thermal cycler ramp speed on amplification of seven target genes using PrimeSTAR GXL enzyme in a 2St PCR**.

**A.** Seven gene targets are readily amplified by adjusting the thermal cycler to a slow ramp speed (2 degrees/s). B. Only one target out of the seven (*Mb0545*) showed an amplicon in a full-speed thermal cycler (6 degrees/s). P. ctrl1 and P. ctrl2 and represent positive controls 1 and 2, *mpb83* amplified using 2 St PCR with an added enhancer at low and full ramp speed, respectively. N. ctrl1 and N. ctrl2 represent negative controls 1 and 2, *Mb0129* amplified using 2St PCR without an enhancer at low and full ramp speed respectively.

**S1 Table. The seven genes used in PCR in S2 Fig, the gene-lengths and GC content**.

